# A critical role of VEGFR2 in lymphatic tumor metastasis

**DOI:** 10.1101/2023.02.16.528814

**Authors:** Taotao Li, Xudong Cao, Fei Zhou, Jing Cui, Beibei Xu, Xiujuan Li, Lena Claesson-Welsh, Taija Makinen, Yulong He

**Affiliations:** Cyrus Tang Hematology Center, Collaborative Innovation Center of Hematology, National Clinical Research Center for Hematologic Diseases, State Key Laboratory of Radiation Medicine and Protection, Cam-Su Genomic Resource Center, Suzhou Medical College of Soochow University, China; Rudbeck, Beijer and SciLifeLab Laboratories, Department of Immunology, Genetics and Pathology, Uppsala University, Uppsala, Sweden; Department of Immunology, Genetics and Pathology, Uppsala University, Uppsala, Sweden

**Keywords:** VEGFR2, tumor lymphangiogenesis, lymphatic metastasis, lymphatic valve, endothelial cell-specific knockout

## Abstract

Vascular endothelial growth factor receptor-2 (VEGFR2) transduces crucial signals for blood vessel growth but its role in the lymphatic system remains incompletely elucidated. By employing genetic mouse models targeting *Vegfr2* in either pan-endothelial cells (ECs) or lymphatic endothelial cells (LECs), we examined roles of VEGFR2 in lymphangiogenesis and in tumor progression. VEGFR2 was differentially expressed in the murine lymphatic system and particularly marked in valves of collecting vessels. The pan-endothelial *Vegfr2* deletion (*Vegfr2^iECKO^*) reduced the dermal lymphatic growth, and a significant decrease in lymphatic valves of pre-collectors was observed in mice with the LEC-specific attenuation of VEGFR2 (*Vegfr2^iLECKO^*). Furthermore, while the primary growth of subcutaneously implanted Lewis lung carcinoma was unaffected in the *Vegfr2^iLECKO^* mouse model, the tumor metastasis to sentinel lymph nodes was efficiently suppressed. In accordance, the tumor-associated lymphangiogenesis was decreased in the *Vegfr2^iLECKO^* mice compared with the control. Findings from this study imply that the lymphatic VEGFR2 regulates valve morphogenesis and promotes lymph node metastasis by regulating the tumor-associated lymphatic formation.

## Introduction

Distant tumor metastasis occurs via the lymphatic and blood vascular systems. Lymphatic formation is governed by a plethora of organotypic lymphangiogenic factors during development and in progression of various diseases [1–5]. Tumor as well as tumor-associated stromal cells secrete lymphangiogenic factors, which are actively involved in the regulation of tumor lymphangiogenesis [6–8]. Suppression of tumor-associated lymphangiogenesis blocks lymphatic tumor metastasis, but does not produce an obvious effect on tumor growth [9].

Vascular endothelial growth factor family members (VEGFs) and their receptors are among the most prominent regulators of angiogenesis and lymphangiogenesis [10–15]. VEGFC/VEGFR3-mediated pathways are crucial for the formation of lymphatic networks and insufficient VEGFR3 signals lead to lymphatic dysplasia and lymphedema [14, 15]. Lymphatic endothelial cells (LECs) also express VEGFR2 (also named KDR or FLK1) [16, 17]. VEGFA as well as the fully processed active forms of VEGFC and VEGFD (VEGFC/D) can activate both VEGFR2 and VEGFR3. VEGFR2 and VEGFR3 also forms heterodimers and respond to VEGFA and VEGFC/D [17–20]. In spite of the biochemical evidence, there is still limited understanding about the role of VEGFR2 in lymphangiogenesis. In zebrafish, VEGFR2 is required for lymphatic development in an organotypic manner [21]. The soluble form of VEGFR2 has been demonstrated to be an endogenous inhibitor of lymphangiogenesis in the cornea [22]. Deletion of *Vegfr2* in endothelial cells using Lyve1-Cre transgenic mice was shown to suppress dermal lymphatic vessel growth [23]. It has also been proposed that VEGFR2 signaling participates in the regulation of lacteal EC junctional integrity as shown in *Vegfr1* (*Flt1*) deleted mice with an increased VEGFA bioavailability [24]. In addition, recombinant adenovirus-mediated overexpression of VEGFA or VEGFE (a VEGFR2-specific factor) induced abnormal lymphangiogenesis [25, 26]. It is worth noting that over-activation of VEGFR2 by excess VEGFA could induce blood vascular leakage and in turn affect the integrity of lymphatic system. Suppression of the VEGFA/VEGFR2-induced vascular leakage and interstitial edema may affect lymphangiogenesis and lymphatic dissemination [27].

To explore the role of lymphatic VEGFR2 in lymphangiogenesis and tumor progression, we employed genetic mouse models targeting *Vegfr2* in either pan-endothelial cells (ECs) or specifically in lymphatic ECs. We found that an efficient deletion of *Vegfr2* could be achieved by using Cdh5-CreERT2 [28], which led to the suppression of lymphangiogenesis in the skin of neonatal mice. Interestingly, we observed that VEGFR2 was highly expressed in lymphatic valves and that the LEC-specific deletion of *Vegfr2 (Vegfr2^iLECKO^*) resulted in a significant decrease in lymphatic valves of pre-collectors. Consistently, attenuation of lymphatic VEGFR2 resulted in a decrease in tumor-associated lymphangiogenesis and significantly suppressed lymphatic tumor metastasis. This suggests that lymphatic VEGFR2 regulates valve morphogenesis of collecting lymphatics and promotes tumor dissemination by regulating tumor-associated lymphangiogenesis.

## Materials and Methods

### Animal models

All animal experiments were performed in accordance with the institutional guidelines of the Soochow University Animal Center. To generate mice with cell-specific *Vegfr2* gene deletion, *Vegfr2^Flox/Flox^* mice [29] were crossed with transgenic mice expressing Cre recombinase in pan-endothelial cells (*Cdh5-CreERT2*) [28] or lymphatic endothelial cells (*Prox1-CreERT2*) [30]. The floxed mice used in this study were maintained on the C57BL/6J background with at least five backcrosses. *Vegfr2^Flox/Flox^;Rosa26^RFP/RFP^* mice was generated as previously reported and expressed the red-fluorescence protein tdTomato after Cre-mediated recombination [31]. In all the phenotype analysis, littermates were used as control.

### Induction of gene deletion

Induction of gene deletion was performed by the systemic administration of tamoxifen. Briefly, new-born pups, including the *Vegfr2^Flox/Flox^;Rosa26^RFP/+^;Cdh5-CreERT2 (Vegfr2^iECKO^*) or *Vegfr2^Flox/Flox^;Rosa26^RFP/+^;Prox1-CreERT2 (Vegfr2^iLECKO^*) and littermate control mice (*Vegfr2^Flox/Flox^;Rosa26^RFP/+^*) were treated with tamoxifen (60 μg per day) by four daily intragastric injections (postnatal days 1-4, P1-4). Adult mice were treated with tamoxifen (1 mg per day) by seven daily intraperitoneal injections. Mice were analyzed at different time points as detailed in the respective figures.

### Xenotransplantation, tumor excision, and analysis

Luciferase tagged Lewis lung carcinoma cells (LLC/Luc) were established by transducing cells with recombinant lentiviral vectors expressing firefly luciferase, and tumor cells with stable expression of luciferase were selected by limiting dilution. Approximately two million tumor cells were subcutaneously implanted into the right abdominal wall of male or female mice (*Vegfr2^Flox/Flox^;Rosa26^RFP/+^;Prox1-CreERT2* or control; 8–12 week old, one tumor per mouse). Tumor growth was monitored by measuring the tumor size every three days for three weeks. In another study, primary tumors were excised two weeks after the tumor implantation, and mice were allowed to recover and analyzed two weeks later. Internal organs including the lungs and axillary lymph nodes were collected for the analysis of tumor metastasis and then further processed for histology.

### In vivo imaging of tumor metastasis and quantification of bioluminescence signal

In vivo imaging was performed using the IVIS Imaging System (Xenogen) as previously described [32]. After the imaging, the animals were euthanized, and organs of interest were collected and imaged *ex vivo*. A region of interest was manually selected, kept constant for all samples and quantified as photons per second using the manufacturer’s imaging software.

### Immunohistochemical staining

For the whole-mount immunostaining, tissues were harvested, fixed in 4 % paraformaldehyde (PFA), blocked with 3 % (w/v) milk in PBS-TX (0.3 % Triton X-100), and incubated with primary antibodies overnight at 4 °C. For the staining of frozen sections, tissues were collected and fixed in 4 % PFA for 2 hours at 4 °C, incubated in 20 % sucrose overnight and then embedded in OCT. Sections were incubated with antibodies against PECAM1 (BD, 553370, 1:500), LYVE1 (Abcam, ab14917, 1:1000), PROX1 (R&D, AF2727, 1:500) and VEGFR2 (R&D, AF644, 1:300). Alexa488- (Invitrogen, A21206 and A21208), Cy5-conjugated secondary antibodies (Jackson ImmunoResearch, 712-175-153, 711-175-152 and 705-175-147) were used for staining. Slides were mounted with 50 % glycerol and analyzed with the Olympus FluoView 3000 confocal microscope.

### Statistical analysis

For the 2-group comparison, the unpaired t test was performed if data passed the D’Agostino-Pearson normality test; otherwise the unpaired nonparametric Mann-Whitney test was applied. For the comparison of the rate of tumor metastasis between two groups, the Chi-square test was performed. All statistical tests were 2-sided using GraphPad Prism 8.

## Results

### Lymphatic endothelial VEGFR2 regulates developmental lymphatic growth and valve formation

To investigate the role of VEGFR2 in lymphatic development, we first employed the inducible pan-endothelial Cre deleter (Cdh5-CreERT2) for the generation of triple transgenic mice (*Vegfr2^Flox/Flox^;Rosa26^RFP/+^;Cdh5-CreERT2*, named *Vegfr2^iECKO^*). Neonatal mice were treated with tamoxifen via the intragastric injection during postnatal days 1 to 4 (Tam1, denoting the first day of tamoxifen treatment) and analyzed at Tam7 (Supplemental Fig. 1A). The expression of reporter RFP indicated the Cre-mediated recombination in endothelial cells and the efficiency of *Vegfr2* deletion was also confirmed by the VEGFR2 immunostaining (Supplemental Fig. 1C). As expected, the induced deletion of pan-endothelial *Vegfr2* (*Vegfr2^iECKO^*) resulted in a dramatic decrease in blood vessel density in the retina, skin, intestinal villi and trachea (Supplemental Fig. 1B-D, F and H). In contrast, the lymphatic phenotypes differed among tissues in the *Vegfr2^iECKO^* mice. While there was a significant decrease in dermal lymphatic density upon the pan-EC deletion of *Vegfr2* (Supplemental Fig. 1C and E), lymphatic formation was not obviously affected in intestinal villi and trachea of the *Vegfr2^iECKO^* mice compared with that of the controls (Supplemental Fig. 1F-H). Consistently, we found in this study that VEGFR2 was differentially expressed in lymphatics among tissues. While VEGFR2 was readily detected in dermal lymphatics, it was relatively low in lacteals and trachea lymphatic vessels (Fig. 1 and Supplemental Fig. 2A-C).

**Fig. 1.**
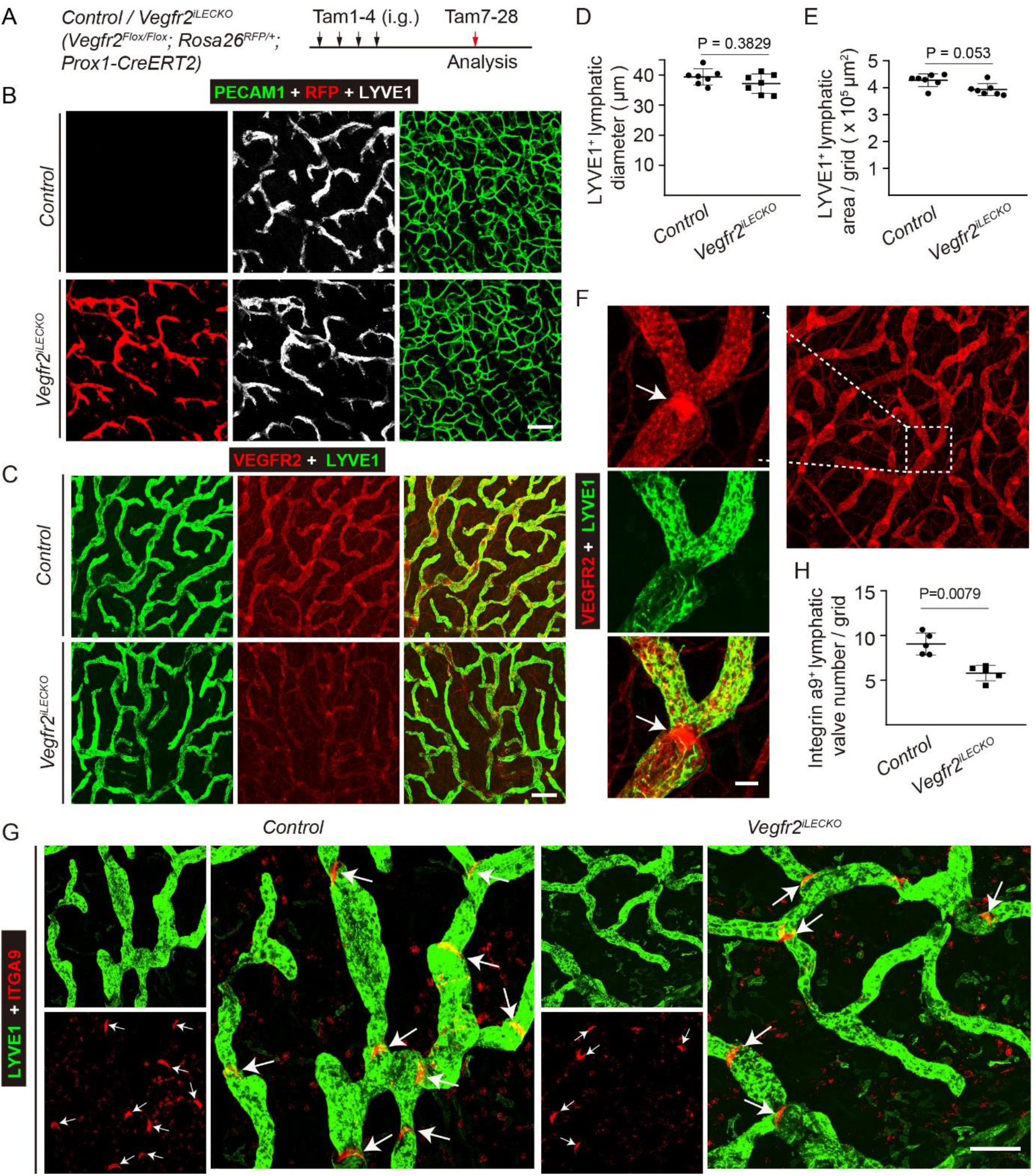
Effect of the LEC-specific deletion of *Vegfr2* on developmental lymphatic growth and valve formation. **A.** Tamoxifen intragastric (i.g.) administration and analysis scheme. **B.** Immunostaining of the abdominal skin from the *Vegfr2^iLECKO^* and control mice (P7) for LYVE1 (grey) and PECAM1 (green). Note that the red fluorescent protein (RFP) signals indicate the expression of reporter gene (tdTomato) after the Cre (*Prox1-CreERT2*)-mediated recombination in lymphatic vessels of the *Vegfr2^iLECKO^* mice. **C.** Analysis of *Vegfr2* deletion of lymphatic vessels in the ventral side of ear skin from the *Vegfr2^iLECKO^* and control mice (P28) by the immunostaining for VEGFR2 (Red) and LYVE1 (green). **D-E.** Quantification of lymphatic vessel diameter was shown in D (LYVE1^+^, μm; Control: 39.34 ± 2.68, n = 7; *Vegfr2^iLECKO^*: 37.10 ± 3.25, n = 7, P = 0.3829). Lymphatic vessel area was quantified as shown in E (LYVE1^+^, × 10^5^ μm^2^ / grid; Control: 4.27 ± 0.24, n = 7; *Vegfr2^iLECKO^*: 3.93 ± 0.22, n = 7, P = 0.053). **F.** Immunostaining of lymphatic vessels in the ear skin for VEGFR2 (red) and LYVE1 (green) from wild-type mice (P21). Arrows indicate the high expression of VEGFR2 in the lymphatic valves of pre-collectors. **G-H.** Analysis of lymphatic valves in the ventral side of ear skin from the *Vegfr2^iLECKO^* and control mice (P28) by the immunostaining for Integrin α9 (Red) and LYVE1 (green). Arrows indicate lymphatic valves. The number of lymphatic valves was quantified as shown in H (Integrin α9^+^, per grid, Control: 9.05 ± 1.23, n = 5; *Vegfr2^iLECKO^*: 5.79 ± 0.87, n = 5, P = 0.0079). Scale bar: 100 μm in B and G; 200 μm in C; 30 μm in F.

To further examine the functions of lymphatic VEGFR2 in development, we used the inducible prospero homeobox 1 (*Prox1*) promoter-driven Cre deleter (*Prox1-CreERT2*) to generate the *Vegfr2^iLECKO^* strain (*Vegfr2^Flox/Flox^;Rosa26^RFP/+^;Prox1-CreERT2*). PROX1 is expressed by various cell types, but is mainly expressed by lymphatic endothelial cells (LECs) in the vascular system [33, 34]. Shown in Fig. 1A is the scheme for the induction of gene deletion. The expression of reporter RFP in LYVE1-postivie vessels was shown in Fig. 1B. Consistent with the observations in *Vegfr2^iECKO^* mice, the loss of LEC-expressed VEGFR2 led to a decrease of lymphatic density, with also a trend of decrease in lymphatic diameter as shown in Fig. 1C-E. Of note, there was still VEGFR2 remaining in the dermal lymphatics of *Vegfr2^iLECKO^* mice as evidenced by the immunostaining analysis (Fig. 1C). The incomplete deletion of *Vegfr2* in the *Vegfr2^iLECKO^* mice may account for the relatively weak lymphatic phenotypes in comparison with that of *Vegfr2^iECKO^* mice (Supplemental Fig. 1). Interestingly, we found that VEGFR2 was highly expressed by lymphatic valves (Fig. 1F). Loss of lymphatic VEGFR2 (*Vegfr2^iLECKO^*) led to a significant decrease in lymphatic valves of pre-collectors (Fig. 1G and H), suggesting that it has a critical role in the regulation of lymphatic valvulogenesis.

### Lymphatic endothelial cell-expressing VEGFR2 in tumor progression

To assess the role of LEC-expressed VEGFR2 in tumor growth and metastasis, the *Vegfr2^iLECKO^* mice were challenged with luciferase-expressing Lewis lung carcinoma (LLC/Luc) implanted subcutaneously. The LEC-specific deletion of *Vegfr2* was induced by tamoxifen treatment (Tam1-7) followed by the implantation of LLC/Luc two days later (Tam3). The scheme for the gene deletion and tumor experiments was as shown in Fig. 2A. Tumor growth was monitored by measuring the tumor size for three weeks (Fig. 2B and Table 1), and tumor weight was also measured when tumor-bearing mice were sacrificed for the analysis at Tam24 (Fig. 2C). This indicates that the LEC-specific *Vegfr2* deletion had no obvious effect on the growth of subcutaneously implanted LLC/Luc tumors as evidenced by the tumor growth curve and tumor weight.

**Fig. 2.**
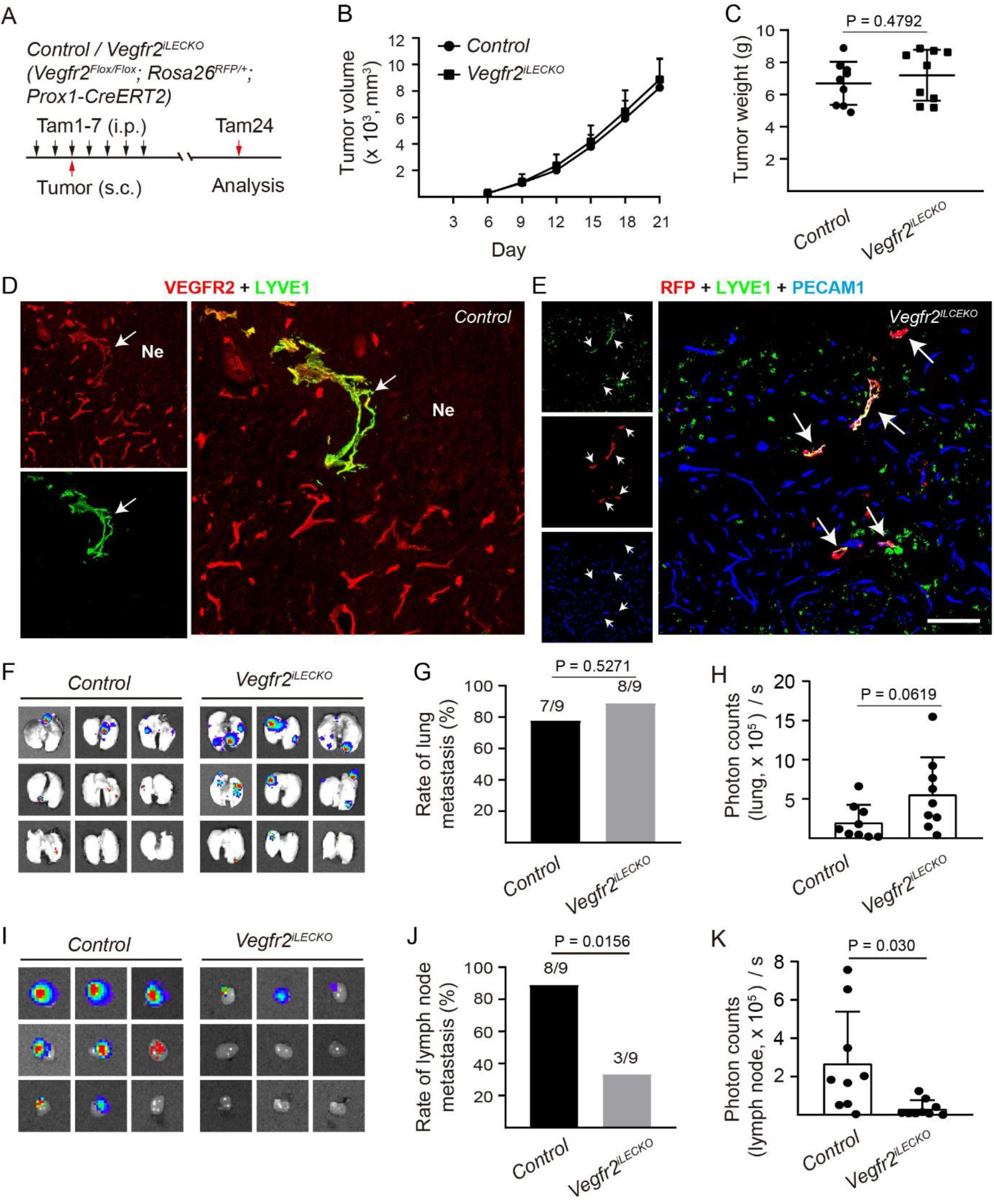
Role of lymphatic endothelial cell-expressing VEGFR2 in tumor progression. **A**. Scheme of tamoxifen administration, tumor implantation and mice analysis. **B-C**. Tumor growth curve (B), and tumor weight at the week 3 stage (LLC/Luc, Tam24) from the *Vegfr2^iLECKO^* and control mice (C; Control: 6.69 ± 1.34 g, *Vegfr2^iLECKO^*: 7.19 ± 1.58 g, n = 9 for each group, P = 0.4792). **D-E.** Immunostaining analysis of VEGFR2 in tumor-associated lymphatic vessels (arrows, D). The reporter RFP indicates the Cre-mediated recombination in tumor lymphatics (arrows, E). **F**. Images of lungs from the LLC/Luc tumor bearing *Vegfr2^iLECKO^* and control mice at Tam24. **G-H**. Rate of lung metastasis (G; Control: 7/9; *Vegfr2^iLECKO^*: 8/9; P = 0.5271), and quantification of bioluminescent signals from the lungs (H; photons/sec × 10^5^, mean ± SD; Control: 2.03 ± 2.21, n = 9; *Vegfr2^iLECKO^*: 5.61 ± 4.70, n = 9; P = 0.0619). **I**. Images of axillary lymph nodes from the LLC/Luc tumor bearing *Vegfr2^iLECKO^* and control mice at Tam24. **J-K.** Rate of lymph node metastasis (J; Control: 8/9; *Vegfr2^iLECKO^*: 3/9; P = 0.0156), and quantification of bioluminescent signals from the axillary lymph nodes (K; photons/sec × 10^5^, mean ± SD; Control: 2.69± 2.69, n=9; *Vegfr2^iLECKO^*: 0.33 ± 0.43, n = 9; P = 0.0300).

**Table 1.**
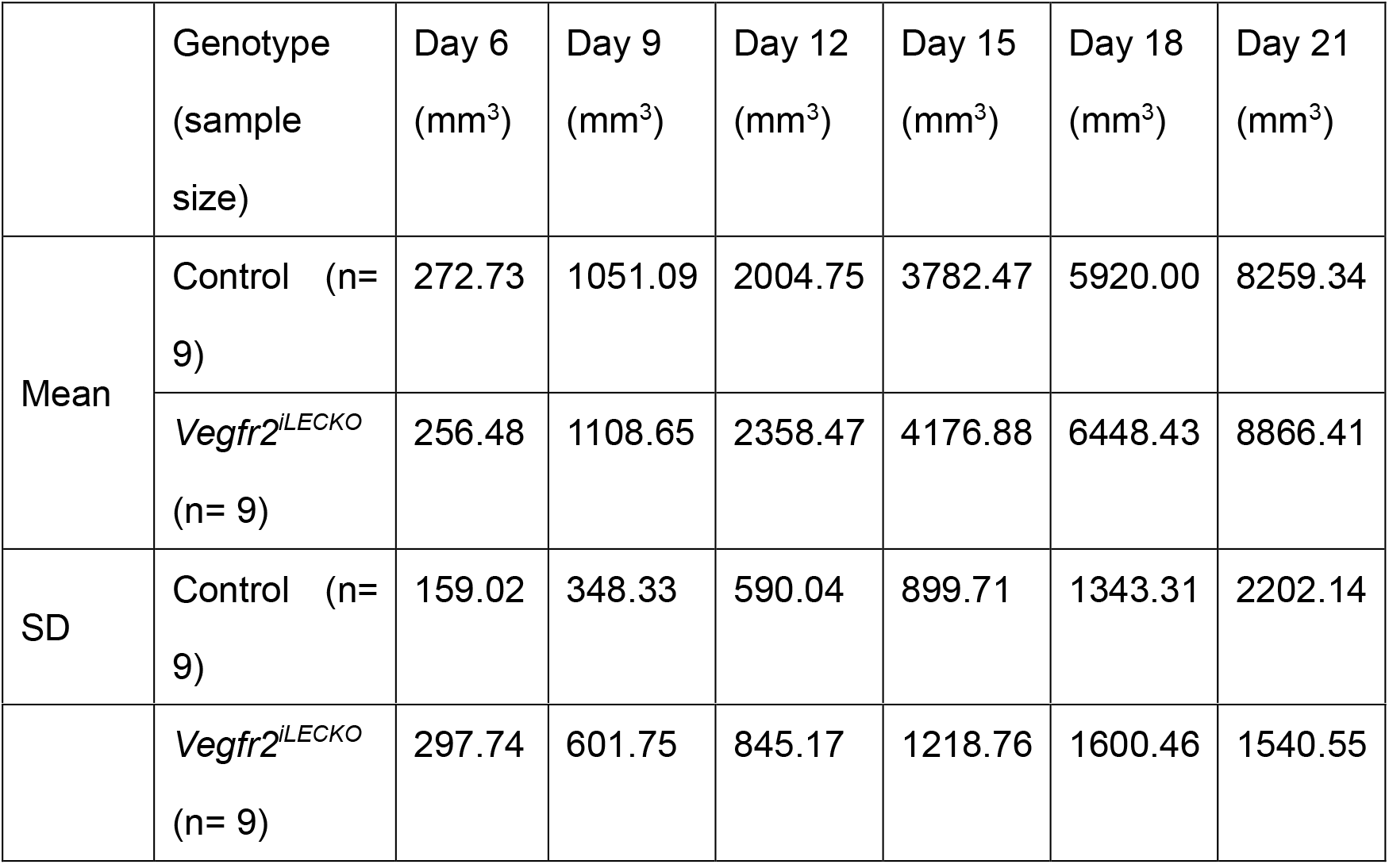
Quantification of subcutaneously implanted tumor volume from the *Vegfr2^iLECKO^* and control mice.

The expression of VEGFR2 was detected in tumor lymphatic vessels (Fig. 2D), and the Cre-mediated recombination was shown by the expression of reporter RFP in LYVE1-positive lymphatics (Fig. 2E). We further assessed the effect of LEC-specific VEGFR2 attenuation on hematogenous and lymphatic tumor metastasis. Shown in Fig. 2F are representative images of lungs from the tumor-bearing *Vegfr2^iLECKO^* and control mice. The rate of lung metastasis (Fig. 2G) and bioluminescent signals from lungs (Fig. 2H) were unaffected after the loss of VEGFR2 expression in the tumor lymphatics. In contrast, lymphatic metastasis to axillary lymph nodes was efficiently reduced in *Vegfr2^iLECKO^* mice (Fig. 2I-K). Shown in Fig. 2I are representative images of lymph nodes from tumor-bearing *Vegfr2^iLECKO^* and control mice. The extent of lymph node metastasis (Fig. 2J) and the bioluminescent signals from lymph nodes (Fig. 2K) showed that loss of VEGFR2 in lymphatic endothelial cells significantly suppressed lymph node metastasis.

### Suppression of tumor-associated lymphangiogenesis by the LEC-specific deletion of *Vegfr2*

To further verify the role of lymphatic VEGFR2 in tumor metastasis, primary tumors (LLC/Luc) were surgically removed after the initial implantation in *Vegfr2^iLECKO^* and control mice. Thereby, the metastatic spread could be examined without the burden of a very large primary tumor. Tumor excision was performed when the primary tumors had grown for two weeks, and the experimental scheme is shown in Fig. 3A. Tumor metastases were analyzed two weeks after the tumor excision. Consistently, loss of VEGFR2 in lymphatic endothelial cells suppressed lymphatic metastasis but had no obvious effect on tumor growth (Fig. 3B) or on the hematogenous tumor metastasis to lungs. Shown in Fig. 3C are representative images of lungs from the tumor-bearing control and *Vegfr2^iLECKO^* mice. The lung metastasis occurred in all the mice from both groups as shown in Fig. 3D, and bioluminescent signals from lungs were quantified (Fig. 3E). Consistent with the above observations, lymph node metastasis was inhibited in *Vegfr2^iLECKO^* mice compared with the control. Shown in Fig. 3F are representative images of lymph nodes from tumor-bearing *Vegfr2^iLECKO^* and control mice. The occurrence of axillary lymph node metastasis was significantly reduced (Fig. 3G), and bioluminescent signals from lymph nodes confirmed the observation (Fig. 3H).

**Fig. 3.**
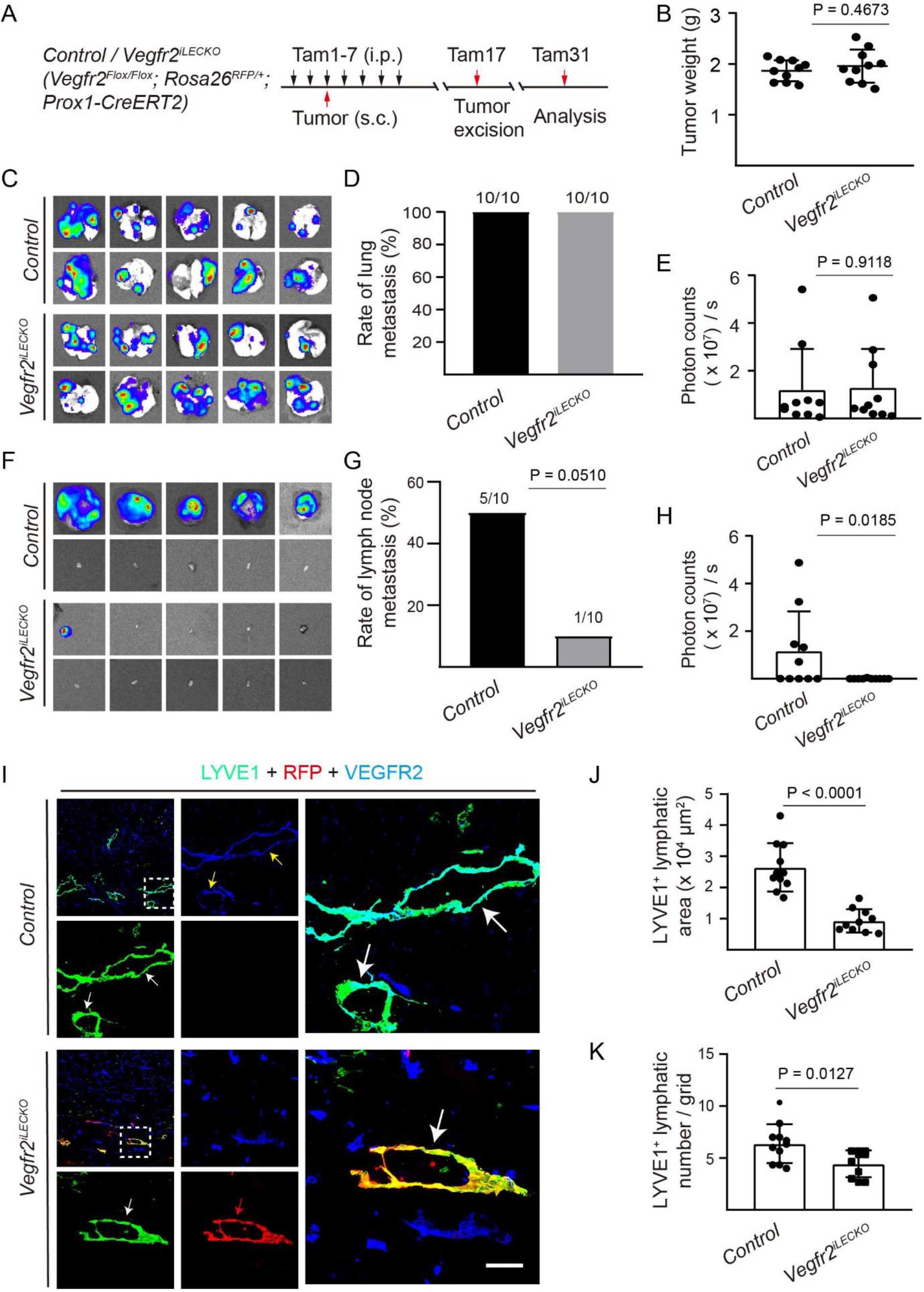
Suppression of tumor-associated lymphangiogenesis and lymph node metastasis by the VEGFR2 attenuation. **A**. Scheme of tamoxifen administration, tumor implantation, tumor excision and mice analysis. **B**. Tumor weight at the week 2 stage (LLC/Luc, Tam17) from the *Vegfr2^iLECKO^* and control mice (Control: 1.87 ± 0.21 g, *Vegfr2^iLECKO^*: 1.96 ± 0.33 g, n = 10 for each group, P = 0.4673). **C-E**. Images of lungs (C, Tam31) from the *Vegfr2^iLECKO^* and control mice with the primary tumors excised at the stage of two weeks (Tam17) after the initial xenograft transplantation. Rate of lung metastasis was shown in D (*Control*: 10/10, and *Vegfr2^iLECKO^*: 10/10), and quantification of bioluminescent signals from the lungs was shown in E (photons/sec × 10^5^, mean ± SD; *Control*: 118.89 ± 172.42, n=10; *Vegfr2^iLECKO^*: 128.34 ± 162.81, n = 10; P = 0.9118). **F-H**. Images of axillary lymph nodes (Tam31) from the *Vegfr2^iLECKO^* and control mice with the primary tumors excised at Tam17 (F). Rate of lymph node metastasis was shown in G (*Control*: 5/10, and *Vegfr2^iLECKO^*: 1/10, P = 0.0510) and quantification of bioluminescent signals from the axillary lymph nodes was shown in H (photons/sec × 10^5^, mean ± SD; *Control*: 115.98 ± 167.03, n = 10; *Vegfr2^iLECKO^*: 0.56 ± 1.41, n = 10; P = 0.0185). **I-K**. Analysis of lymphatic vessels in LLC/Luc tumors (Tam17) from the *Vegfr2^iLECKO^* and control mice by the immunostaining for LYVE1 (green), VEGFR2 (blue) and RFP (red). White arrows point to the tumor-associated lymphatic vessels, where VEGFR2 was readily detected in controls (yellow arrows in I). VEGFR2 was not detectable in lymphatics expressing LYVE1 (white arrows in I) and the reporter RFP (red arrows, I) in the *Vegfr2^iLECKO^* mice. Quantification of tumor-associated lymphatic vessel area was shown in J (LYVE1^+^, x 10^4^ μm^2^ / grid; Control: 2.64 ± 0.77, n = 11; *Vegfr2^iLECKO^*: 0.93 ± 0.37, n = 10, P < 0.0001). Quantification of tumor-associated lymphatic vessel number was shown in K (Control: 6.36 ± 1.87 per grid, n = 11; *Vegfr2^iLECKO^*: 4.43 ± 1.30 per grid, n = 10, P = 0.0127). Scale bar: 50 μm in I.

To explore the pathological mechanism underlying the inhibition of lymphatic metastasis after the LEC-specific *Vegfr2* deletion, we analyzed the lymphatic and blood vascular growth in tumors by the immunostaining for LYVE1 and VEGFR2. The LYVE1 positive lymphatic vessels in three microscopic fields of the highest vessel area were quantified using Image-Pro Plus 6.0. There was a significant decrease in lymphatic area, reflecting the lymphatic vessel with lumens, and also vessel numbers in tumor xenografts from the *Vegfr2^iLECKO^* mice compared with that of the control (Fig. 3I-K). The deletion of *Vegfr2* was confirmed by the immunostaining for VEGFR2 and LYVE1 (Fig. 3I). The results from tumor and developmental studies suggest that lymphatic VEGFR2 regulates valve morphogenesis and promotes lymph node metastasis by regulating the tumor-associated lymphatic vessel growth. VEGFR2 may act independently or interact with VEGFR3 by forming VEGFR2/3 heterodimers in the regulation of lymphangiogenesis (Fig. 4).

**Fig. 4.**
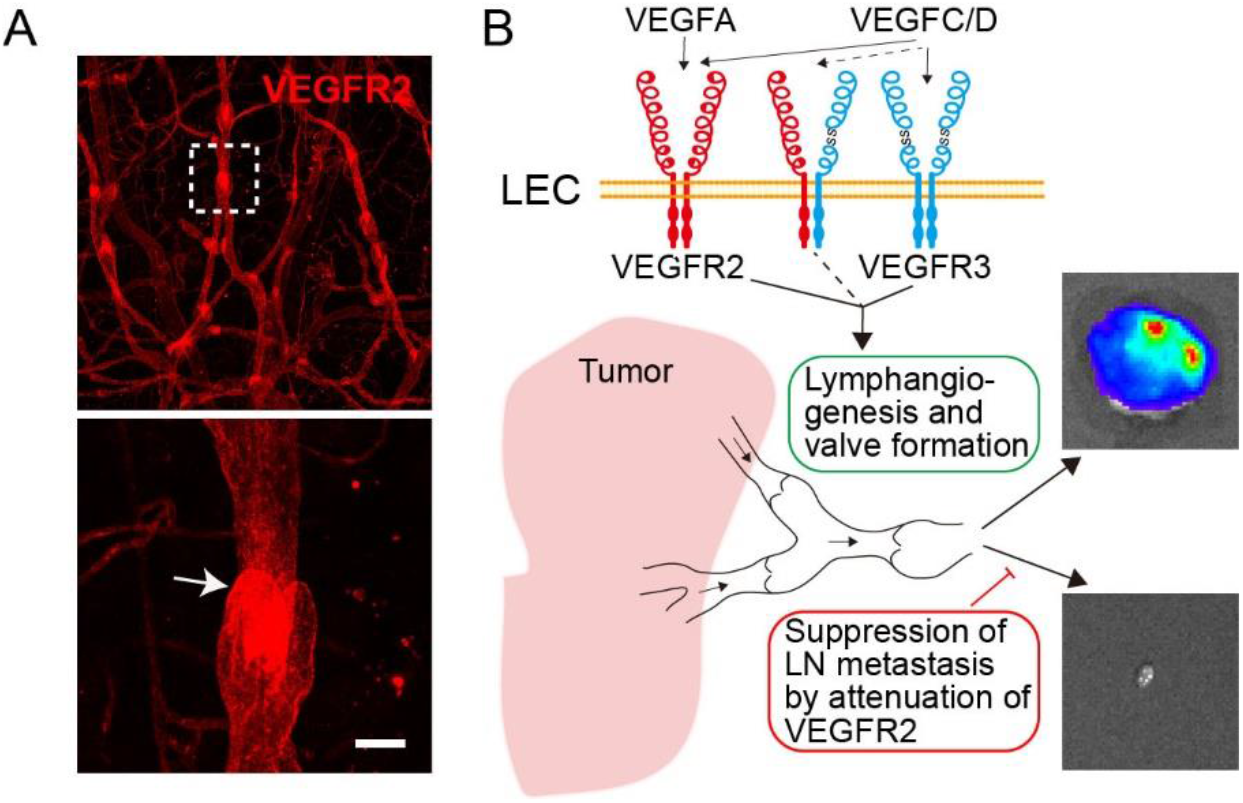
A proposed model of VEGFR2 in lymphatic tumor metastasis. **A.** VEGFR2 is highly expressed in lymphatic valves (arrow) of collecting lymphatics as shown in the dorsal side of ear skin (arrows). **B**. It has been well established that VEGFR3 is essential for lymphangiogenesis and lymphatic tumor metastasis [9, 48]. Interestingly, findings from this study show that attenuation of VEGFR2 produces a dramatic inhibition of lymphatic tumor metastasis. We propose that VEGFR2 expressed by lymphatic endothelial cells (LECs) functions either independently or together with VEGFR3 by forming VEGFR2/3 heterodimers in the regulation of tumor-associated lymphatic formation and maturation, which facilitates the lymph node (LN) metastasis. Scale bar: 30 μm in A.

## Discussion

We show in this study that the induced attenuation of VEGFR2 in lymphatic endothelial cells suppresses lymph node metastasis by reducing the tumor-associated lymphangiogenesis. Consistently, VEGFR2 insufficiency in either pan-endothelial cells or lymphatic endothelial cells decreases dermal lymphatic growth in development while lymphatics in other organs are less affected. This may be due to the organ-specific requirement of VEGFR2 during lymphatic formation [21, 35]. We have also found that VEGFR2 is highly expressed in lymphatic valves and that the LEC-specific deletion of *Vegfr2* results in a significant decrease in lymphatic valves of pre-collectors. These findings support a direct role of VEGFR2 in the lymphatic system at both physiological and pathological conditions.

VEGFR2 is essentially required for angiogenesis in development and tumor growth [10, 13, 36]. However, its role in the lymphatic system is much less understood. It was previously reported that there was a lymphatic hypoplasia in skin after the conditional deletion of *Vegfr2* in endothelial cells using a *Lyve1-Cre* transgenic mouse line [23]. As LYVE1 is also expressed in some blood vascular endothelial cells (BECs) [18, 37], the underdeveloped dermal lymphatic network could be secondary to the suppression of angiogenesis and/or BEC-derived lymphangiogenic factors after the loss of endothelial VEGFR2. We also found in this study that the pan-endothelial VEGFR2 attenuation (*Vegfr2^iECKO^*) led to the decrease of lymphatic growth in skin of neonatal mice. To further investigate the role of LEC-expressed VEGFR2 in lymphatic development, we established a strain with the LEC-specific deletion of *Vegfr2* (*Vegfr2^iLECKO^*). We found that VEGFR2 insufficiency led to the decrease in lymphatic density as well as a significant decrease in lymphatic valves of pre-collectors in skin. Consistently, NRP1, a coreceptor of VEGFR2, regulates the formation of lymphatic valves and mice carrying a mutation in the SEMA3A binding site of NRP1 were shown to display lymphatic valve defects [38, 39]. This suggests that VEGFR2 signaling pathway participates in the regulation of lymphatic valve morphogenesis. However, VEGFR2 attenuation did not produce an obvious effect on lymphatic development in several other tissues examined at the same stage including trachea or intestine villi in this study. An organ-specific function of VEGFR2 was also reported in Zebrafish, where it was shown to be required for the rostral craniofacial lymphangiogenesis but not for the caudal trunk lymphangiogenesis [21]. Detailed mechanisms underlying the organotypic function of VEGFR2 in lymphatic development warrant further investigation although the differential expression of VEGFR2 among tissues could be one reason as shown in this study.

Accumulating evidence also points to a potentially direct role of VEGFA/VEGFR2-mediated signals in lymphangiogenesis in pathological conditions. VEGFA-VEGFR2 signaling has also been shown to induce lymphatic growth in cutaneous leishmaniasis [40]. In a mouse corneal suturing model, the expression of soluble VEGFR2 (sKDR) by the splice-shifting morpholinos or the systemic administration of neutralizing antibodies against VEGFR2 suppressed corneal angiogenesis and lymphangiogenesis [41, 42]. VEGFA and the specific VEGFR3 activator VEGFC156S were shown to induce a similar set of target genes as well as some specific ones in primary human lymphatic endothelial cells [43]. Overexpression of VEGFA or VEGFE via an adenoviral vector induced abnormal lymphangiogenesis [25, 26]. VEGFA was shown to induce lymphangiogenesis via the upregulation of VEGFC in vascular endothelial cells [44]. Furthermore, VEGFC has been shown to promote the VEGFR2/VEGFR3 heterodimerization, which may mediate signals for lymphangiogenesis in cauterised cornea and tumor [17, 45]. It was also reported that VEGFR2 polymorphism (rs2071559, T/C) had an association with lymphatic metastasis in patients with nasopharyngeal carcinoma [46]. Tyrosine kinase inhibitors or neutralizing antibodies against VEGFR2 were reported to have a suppressive effect on tumor-associated lymphatics [47]. We further demonstrated in this study that attenuation of lymphatic VEGFR2 suppressed tumor-associated lymphangiogenesis with a significant inhibition of lymphatic tumor metastasis.

In summary, we have shown in this study that VEGFR2 is required for the dermal lymphatic growth, particularly the valve formation of collecting lymphatic vessels in development. The attenuation of lymphatic VEGFR2 also suppresses the tumor-associated lymphangiogenesis and lymphatic metastasis. Therefore, VEGFR2 could be employed for modulating both angiogenesis and lymphangiogenesis in cancer. Targeting VEGFR2 provides a therapeutic potential for blocking tumor dissemination via both lymphatic and blood vascular routes.

## Acknowledgement

We thank Dr. Janet Rossant for kindly providing us the *Vegfr2^Flox/Flox^* mouse line and Dr. Yoshiaki Kubota of Keio University for the *Cdh5-CreERT2* mouse line, and staff in the Animal facility of Soochow University for technical assistance.

