## Supplemental figures and figure legends for "A critical role of VEGFR2 in lymphatic tumor metastasis"

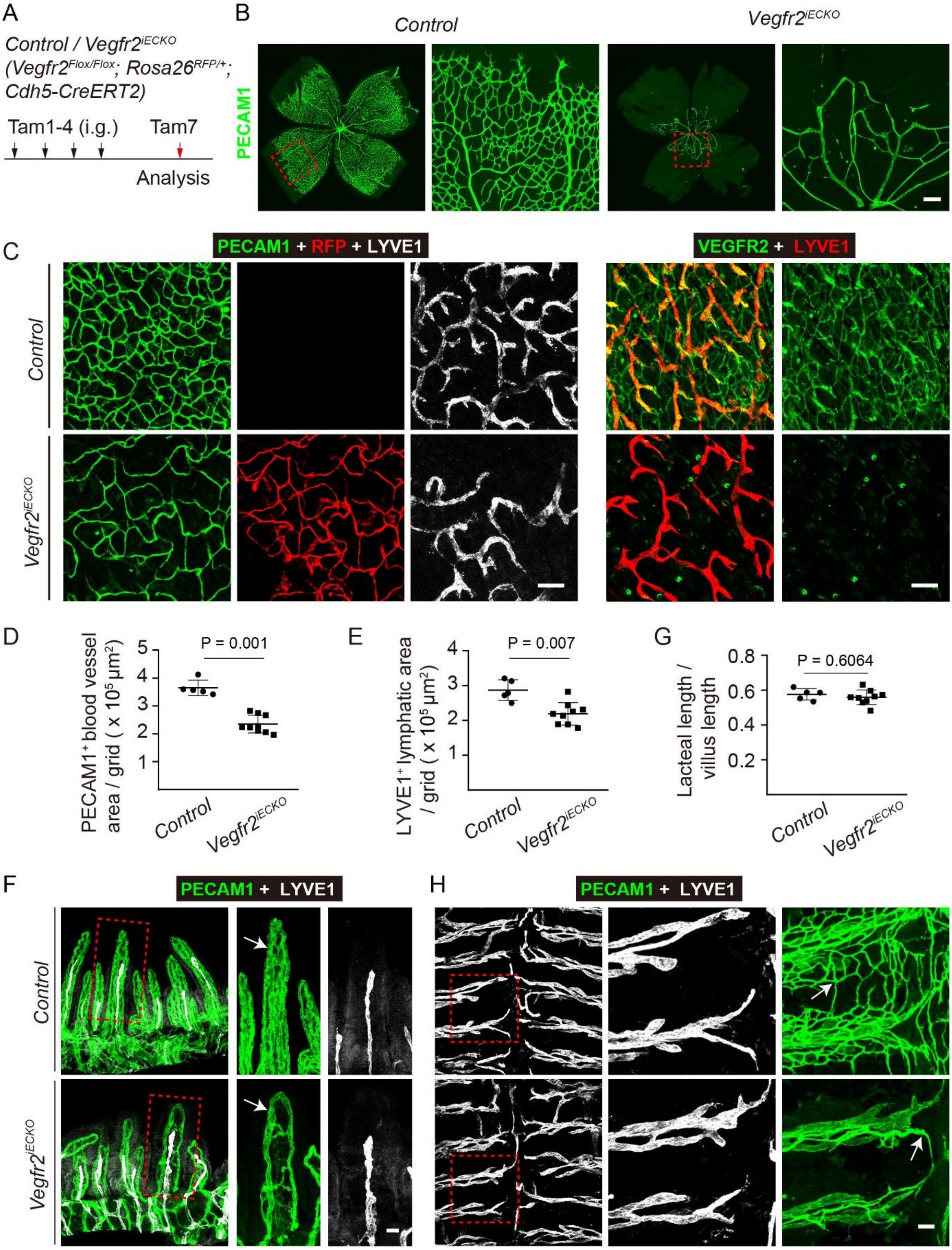


**Supplemental Fig. 1 Analysis of the developmental lymphangiogenesis after the pan-endothelial deletion of *Vegfr2*. A**. Tamoxifen intragastric (i.g.) administration and analysis scheme for *Vegfr2*$^{iECKO}$ and control mice. **B**. Analysis of blood vessels in retina from the

*Vegfr2$^{iECKO}$* and control mice (P7) by the immunostaining for PECAM1 (green). **C.** Left panel: Analysis of blood vessels and lymphatic vessels in the abdominal skin from the *Vegfr2$^{iECKO}$* and control mice (P7) by the immunostaining for PECAM1 (green) and LYVE1 (grey). Right panel: Analysis of *Vegfr2* deletion in abdominal skin from the *Vegfr2$^{iECKO}$* and control mice (P7) by the immunostaining for VEGFR2 (green) and LYVE1 (red). Note that RFP signals indicate the expression of reporter gene (red fluorescent protein) by the Cre-mediated (*Cdh5-CreERT2*) recombination in blood and lymphatic vessels of the *Vegfr2$^{iECKO}$* mice. **D-E**. Quantification of blood vessel area (D, PECAM1$^+$, x $10^5$ μm$^2$ / grid, Control: 3.65 ± 0.28, n = 5; *Vegfr2$^{iECKO}$*: 2.35 ± 0.32, n = 9, P = 0.001) and lymphatic vessel area (E, LYVE1$^+$, x $10^5$ μm$^2$ / grid, Control: 2.87 ± 0.30, n = 5; *Vegfr2$^{iECKO}$*: 2.19 ± 0.32, n = 9, P = 0.007). **F-H.** Analysis of blood vessels and lymphatic vessels in the intestinal villi (F) and trachea (H) from the *Vegfr2$^{iECKO}$* and control mice (P7) by the immunostaining for PECAM1 (green) and LYVE1 (grey). Quantification of the ratio of lacteal to villus length was shown in G (Control: 0.58 ± 0.03, n = 5; *Vegfr2$^{iECKO}$*: 0.56 ± 0.04, n = 9, P = 0.6064). Scale bar: 100 μm in B, C; 30 μm in F; 50 μm in H.

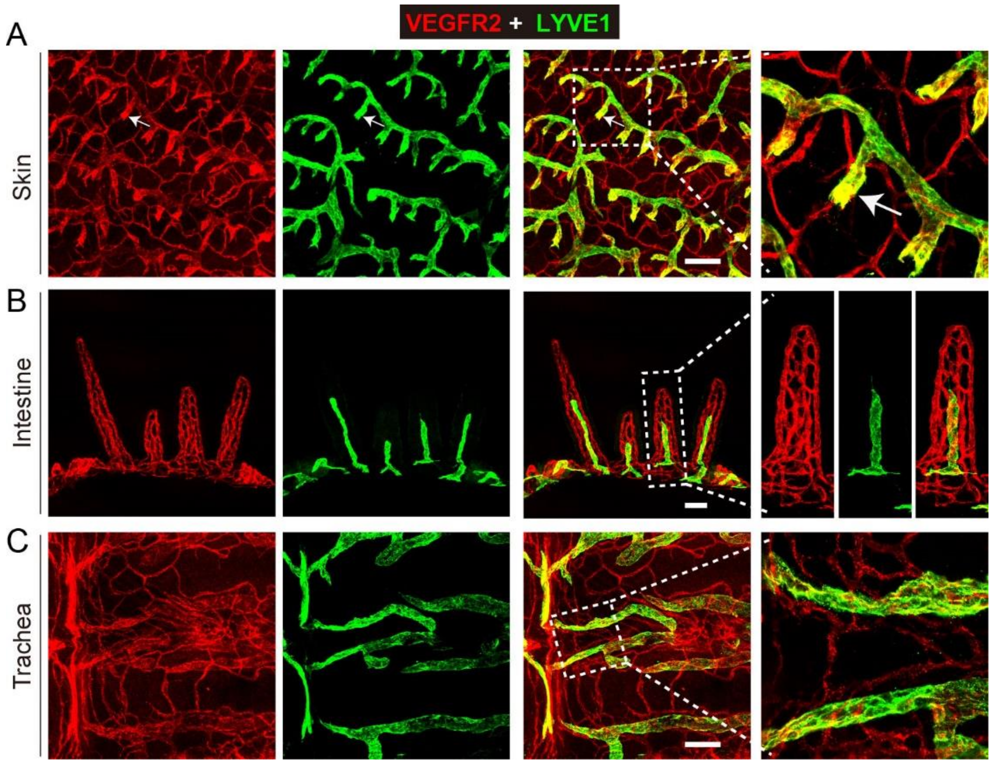


**Supplemental Fig. 2 Analysis of VEGFR2 expression in different tissues. A-C**. Immunostaining of VEGFR2 expression in lymphatic vessels of the ventral skin (A), villi (B) and trachea (C) in wild-type mice (P7; VEGFR2, red; LYVE1, green). Arrows indicate the high expression of VEGFR2 in dermal initial lymphatic vessels. Scale bar: 100 μm in A and C; 200 μm in B.
